# Mdig shapes 3D chromatin architecture of immune checkpoint genes and limits metastasis of triple negative breast cancer cells

**DOI:** 10.64898/2025.12.23.696275

**Authors:** Ziwei Wang, Ziqi Liu, Chitra Thakur, Yiran Qiu, Jun Wang, Mehdi Damaghi, John D. Haley, Fei Chen

## Abstract

Despite decades of progress in metastasis research, the identity of bona fide metastasis-driving genes remains elusive. By shifting the focus from individual genes to the three-dimensional (3D) architecture of the genome, we identify the mineral dust-induced gene (mdig) as a chromatin-topology regulator that suppresses immune-checkpoint activation and metastasis in triple-negative breast cancer (TNBC). Re-expression of mdig in mdig-knockout (KO) cells markedly reduced liver metastasis, whereas mdig loss activated epithelial-mesenchymal transition (EMT), T-cell exhaustion, and antigen-presentation pathways. Across multiple cancer types, high mdig expression predicted improved responses to immune checkpoint blockade (ICB). Mechanistically, mdig depletion derepressed inhibitory checkpoint genes, including PD-L1, PD-L2, LGALS9, and PLXNB1/2, accompanied by increased H3K9me3 and H3K36me3 at these loci. Hi-C profiling revealed that mdig maintains topologically associating domains (TADs) and long-range repressive loops at checkpoint loci; loss of mdig disrupted these structures, enabling coordinated activation of immune-evasion programs. Genome-wide analyses showed extensive TAD remodeling in mdig KO cells, with pronounced alterations on chromosome X. Collectively, these findings position mdig as a key chromatin-architectural regulator of immune evasion and metastatic behavior, underscoring its potential as a biomarker for ICB responsiveness in TNBC and beyond.

**Significance statement:** Metastasis remains the leading cause of cancer mortality, yet bona fide metastasis-driving genes remain difficult to define. By examining three-dimensional genome architecture rather than individual gene function, we identify mdig as a chromatin-topology regulator that restrains immune evasion and metastasis in triple-negative breast cancer. Loss of mdig disrupts TAD structures and long-range repressive loops at immune-checkpoint loci, enabling coordinated activation of EMT, antigen-presentation, and T-cell-exhaustion programs. Restoring mdig expression markedly reduces liver metastasis, and high mdig levels across cancers predict improved responses to immune-checkpoint blockade. These findings establish mdig as a key regulator of chromatin organization and a potential biomarker for immunotherapy responsiveness.

## Introduction

Cancer metastasis research has long been guided by a gene-centric paradigm, in which investigators search for individual “driver” genes or small gene sets as the primary regulators of dissemination and colonization (1). This approach has yielded important insights, identifying transcription factors, signaling molecules, and adhesion regulators that promote invasion, angiogenesis, and immune evasion. However, this framework overlooks a more fundamental layer of gene control: the three-dimensional (3D) organization of the genome (2). Chromatin folding determines the spatial proximity of enhancers, promoters, and insulators, thereby enabling or constraining entire transcriptional programs. Alterations in higher-order chromatin architecture can activate metastatic gene networks or silence metastasis-suppressive pathways without any change in DNA sequence.

Recent studies employing Hi-C, micro-C, and related techniques have revealed that chromatin architecture is frequently remodeled in aggressive tumors (3). Disruption of topologically associating domains (TADs), aberrant formation or loss of long-range enhancer-promoter loops, and compartment switching have all been linked to invasion, acquisition of stem-like traits, and therapy resistance across multiple cancer types. These 3D genome alterations can act as upstream structural drivers, coordinating the simultaneous activation of metastasis-associated gene programs, an effect far broader than that of single-gene mutations (4). Despite these advances, most metastasis studies still rely primarily on transcriptomic or mutational profiling, leaving the structural genome largely unexamined.

We previously demonstrated that the mineral dust-induced gene (mdig) functions as an anti-metastatic factor, and that knockout (KO) of mdig in the triple-negative breast cancer (TNBC) cell line MDA-MB-231 promotes liver and lung metastasis (5, 6). Here, we provide evidence that re-expressing mdig in mdig KO cells substantially reduces liver metastatic burden. Transcriptomic profiling revealed that mdig depletion enforces the expression of genes involved in epithelial-mesenchymal transition (EMT), metastasis, and immune modulation, suggesting that mdig act as a non-canonical tumor suppressor that curtails metastatic progression by regulating tumor immunity. This interpretation is supported by clinical datasets showing that high mdig expression correlates with significantly improved survival in patients receiving immune checkpoint blockade (ICB) therapies, including anti-PD-1, anti-PD-L1, and anti-CTLA-4.

Consistent with this, our data indicate that mdig suppresses major immune suppressive genes, including MHC class I and II genes, *PD-L1*, *PD-L2*, *PLXNB1*, *PLXNB2*, and others. Analysis of 3D chromatin architecture further confirmed that mdig depletion alters TAD organization at genomic loci harboring immune checkpoint and pro-metastasis genes. Together, these findings uncover a previously unexplored architectural mechanism through which mdig regulates immune checkpoint expression and metastatic competence. They also illustrate a conceptual shift from gene-centric models toward a spatial-genome perspective, an essential step toward a more complete, predictive, and therapeutically actionable understanding of metastasis biology.

## Results

### Restoring mdig in mdig KO cells blocks liver metastasis in TNBC

We previously demonstrated that knocking out mdig enhances lung and liver metastasis of MDA-MB-231 cells *in vivo* (5, 7). To determine whether this phenotype reflects true loss of mdig rather than clonal artifacts arising from CRISPR-Cas9 gene editing, we re-expressed mdig in mdig KO cells using a lentiviral vector. Mdig KO cells transduced with either an empty vector (mdig KO-vec) or an mdig-expressing vector (mdig KO-mdig) were orthotopically injected into the 5^th^ mammary fat pad of female NSG mice, and primary tumor progression and metastasis were monitored. Nine weeks after injection, all mice were euthanized for metastatic assessment. Restoring mdig expression had minimal impact on primary tumor growth (data not shown), but markedly reduced liver metastasis. At necropsy, a number of visible liver metastatic lesions was observed in mice injected with mdig KO-vec cells (Fig. 1A, left), whereas no visible liver metastases were detected in any of the six mice injected with mdig KO-mdig cells (Fig. 1A, right). H&E staining further confirmed a dramatic reduction in liver metastatic nodules in the mdig KO-mdig group (Fig. 1B). Re-analysis of RNA-seq data comparing mdig KO and WT cells (shrunken log2FC ≥ 1.0) using MSigDB Cancer Hallmark 2020 and GO Biological Process 2025 revealed enrichment of genes associated with EMT, T-cell exhaustion, MHC-mediated antigen presentation, cell adhesion, and migration (Figs. 1C & 1D). Together, these findings suggest that mdig suppresses EMT and inhibitory tumor immune pathways, thereby restricting liver metastasis in TNBC.

**Figure 1.**
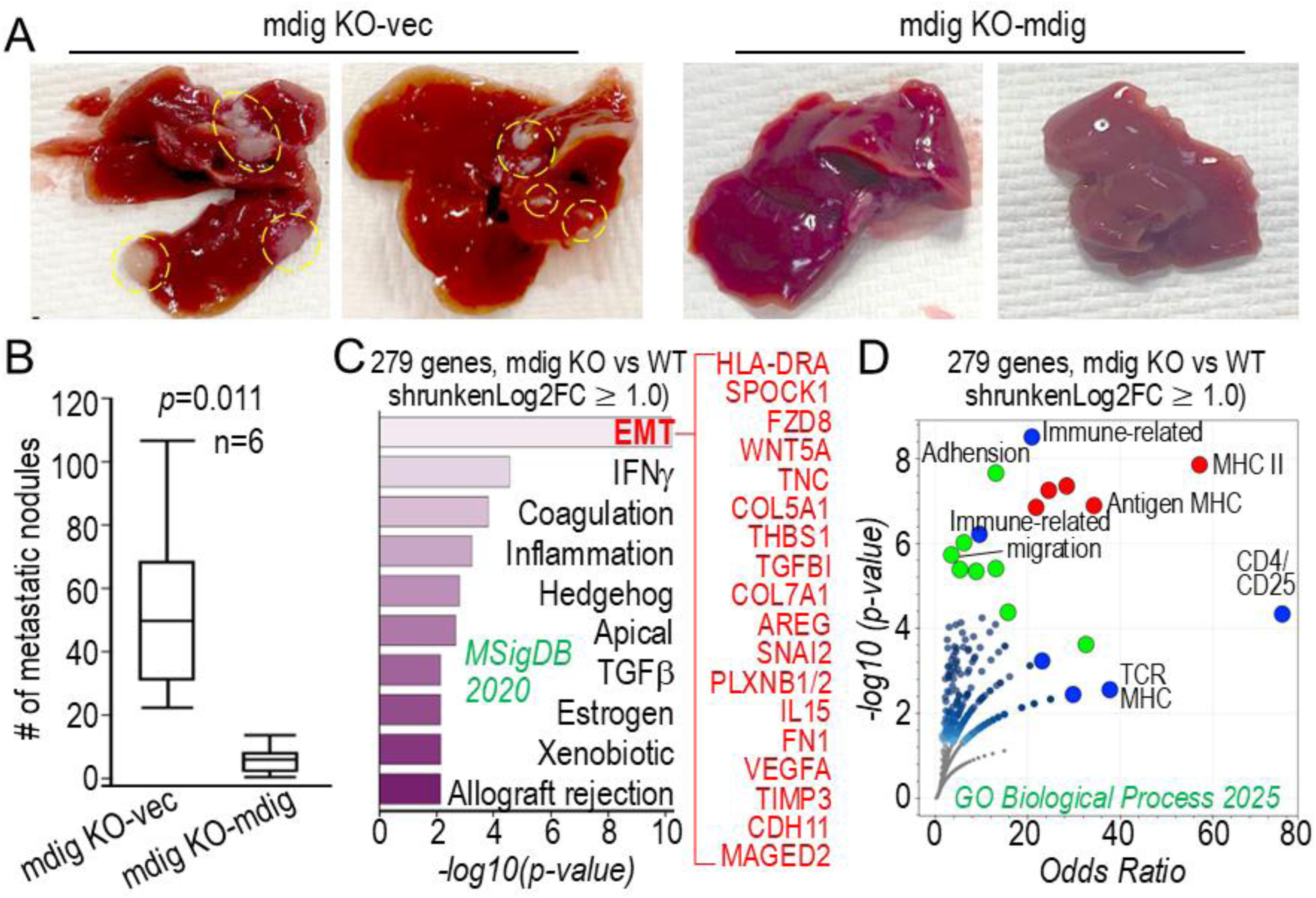
Restoring mdig expression in mdig KO TNBC cells suppresses liver metastasis. **(A)** Representative gross images of livers harvested from NSG mice nine weeks after orthotopic injection of mdig KO-vec or mdig KO-mdig MDA-MB-231 cells. Multiple visible metastatic lesions were present in mice injected with mdig KO-vec cells, whereas no visible liver metastases were detected in mice injected with mdig KO-mdig cells (n = 6 per group). **(B)** Quantification of H&E-stained liver sections showing a significant reduction in metastatic nodules in the mdig KO-mdig group compared with mdig KO-vec controls. **(C)** MSigDB Cancer Hallmark 2020 enrichment analysis of differentially expressed genes (shrunken log2FC ≥ 1.0) identified upregulation of pathways associated with EMT, T-cell exhaustion, antigen presentation, and cell adhesion/migration in mdig KO versus WT cells. **(D)** GO Biological Process 2025 enrichment analysis showing similar enrichment of EMT- and immune-related processes.

### High mdig expression improves the efficacy of immune checkpoint blockade (ICB) therapy

The strong influence of mdig on immune-regulatory pathways in TNBC cells aligns with our earlier observation that heterozygotic knockout of mdig (mdig^+/−^) mice exhibit increased infiltration of Foxp3⁺ regulatory T cells (Tregs) in fibrotic lungs compared with wild-type (WT) mice (8). Given that elevated Tregs undermine the effectiveness of ICB therapy by suppressing CD8^+^ T-cell-mediated tumor surveillance and higher Treg infiltration associated with poorer prognosis of cancer patients (9), we subsequently evaluated whether mdig expression correlates with clinical responses to ICB. Due to the limited availability of ICB response data in breast cancer, we analyzed aggregated survival datasets from patients with bladder cancer, esophageal adenocarcinoma, glioblastoma, hepatocellular carcinoma, melanoma, lung cancer, and urothelial carcinoma. Across these tumor types, high mdig expression was consistently associated with significantly improved progression-free survival (PFS) in patients treated with anti-PD-1, anti-PD-L1, or anti-CTLA-4 therapies (Fig. 2A). This association was further corroborated by higher mdig levels in responders compared with non-responders among patients with metastatic disease treated with anti-PD-1 or anti-CTLA-4 agents (Fig. 2B). These results suggest that mdig may act as a non-canonical tumor suppressor that shapes metastatic progression and modulate response to ICB. To contextualize mdig within known tumor suppressor landscapes, we compared its prognostic performance with that of 35 established tumor suppressor genes. Notably, mdig clustered with *RB1*, *APC*, *NF2*, *WT1*, *TSC1*, *CHEK2*, and *CYLD*, genes whose high expression predicts improved survival across all three major ICB classes. In contrast, several canonical suppressors such as *TP53*, *RAD51*, and *KISS1* were associated with favorable outcomes in only one or two ICB categories (Fig. 2C). This pattern may help explain the reduced mdig expression observed in a subset of breast cancers (Fig. 2C, right). Further supporting a tumor-suppressive function, pathway analysis of 766 genes negatively correlated with mdig expression in breast cancer (Spearman’s Rho ≤ −0.2) revealed strong enrichment for oncogenic programs, including KLF4 and SNAI1/2 signaling, estrogen response, EMT, and related transcriptional networks (Fig. 2D).

**Figure 2.**
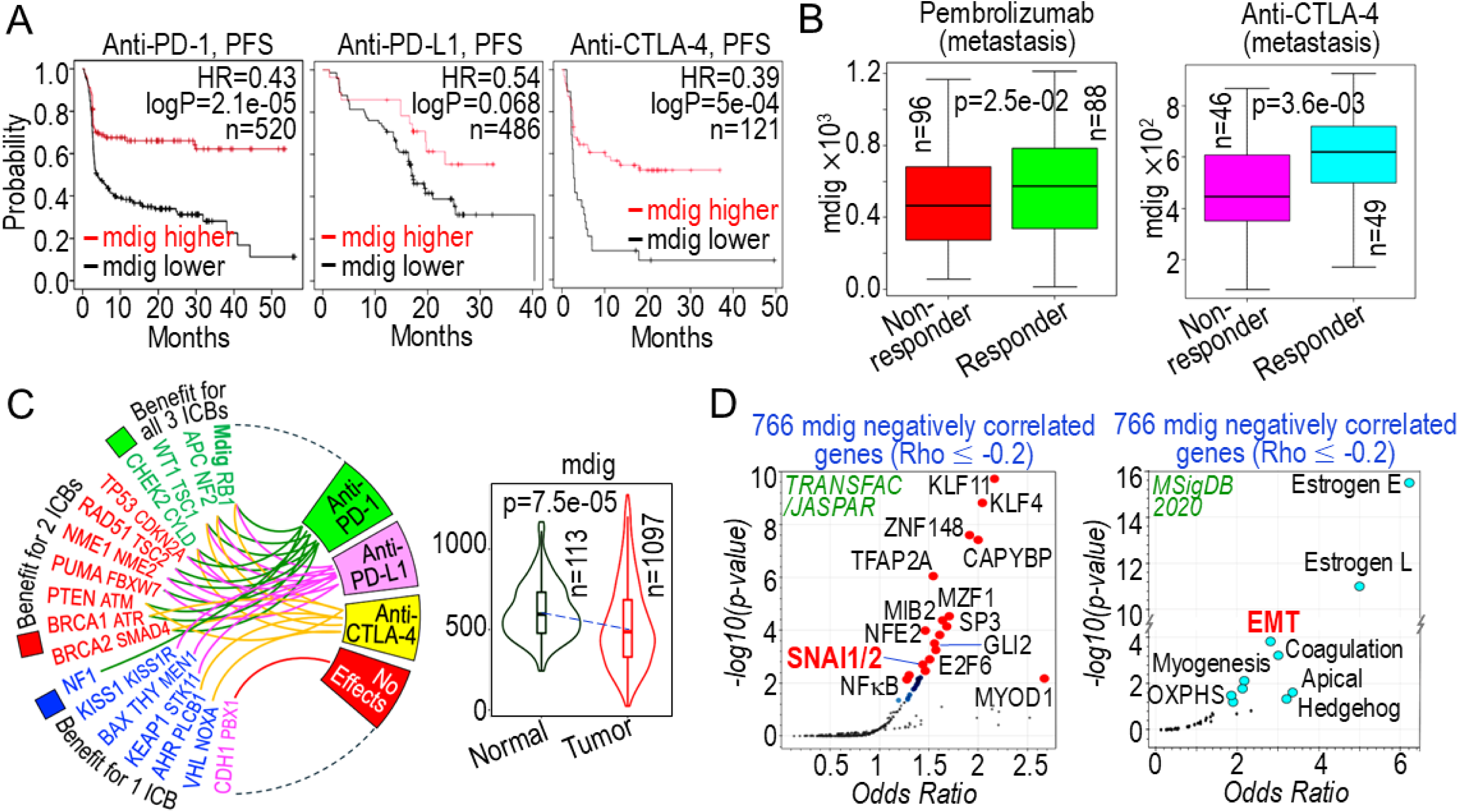
High mdig expression is associated with improved response to ICB therapy. **(A)** Kaplan-Meier analysis of PFS in patients across multiple cancer types treated with anti-PD-1, anti-PD-L1, or anti-CTLA-4 therapies. Patients with high mdig expression exhibited significantly improved PFS compared with those with low mdig expression. **(B)** Comparison of mdig expression levels in responders versus non-responders among patients with metastatic disease treated with anti-PD-1 or anti-CTLA-4 agents, showing higher mdig levels in responders. **(C)** Circos plot showing prognostic performance of mdig relative to 35 established tumor suppressor genes across the three major ICB classes. Mdig clustered with *RB1*, *APC*, *NF2*, *WT1*, *TSC1*, *CHEK2*, and *CYLD*, whose high expression predicts favorable outcomes, whereas canonical suppressors such as *TP53*, *RAD51*, and *KISS1* were associated with favorable outcomes in only one or two ICB categories. Right panel: reduced mdig expression observed in a subset of breast cancers. **(D)** Pathway enrichment analysis of 766 genes negatively correlated with mdig expression in breast cancer (Spearman’s Rho ≤ −0.2), revealing strong association with oncogenic programs, including KLF4 and SNAI1/2 signaling, estrogen response, EMT, and related transcriptional networks.

### Negative regulation of inhibitory immune checkpoint genes by mdig

The strong association between high mdig expression and improved outcomes following ICB therapy suggests that mdig plays a role in modulating tumor immunity. This is consistent with our previous observations of increased Treg infiltration in the lungs of mdig^+/−^ mice exposed to silica particles (8), as well as evidence that mdig interacts with molecules involved in antigen presentation (10). Hyperactivation of immunosuppressive Treg is known to compromise ICB efficacy by suppressing CD8^+^ T cell function (9). To directly assess whether mdig regulates immune checkpoints, we compared the expression of canonical and non-canonical inhibitory checkpoint genes in WT and mdig KO cells. Strikingly, mdig KO cells exhibited marked upregulation of multiple checkpoint genes, including *HLA-DRA/DRB1*, *LGALS9*, *PD-L1*, *PD-L2*, *SELPLG*, *PLXNB1*, and *PLXNB2* (Figs. 3A & 3B). Clinically, PLXNB2 and PD-L1 are elevated in metastatic breast cancer, and mdig expression inversely correlates with PLXNB1 and PLXNB2 in patient tumors (Figs. 3C & 3D). PLXNB1 and PLXNB2 have recently emerged as key regulators of tumor immunity. Plxnb1^-/-^ BALB/c mice show enhanced M1 polarization of tumor-associated macrophages and improved response to anti-PD-1 therapy (11), whereas Plxnb2 deficiency reduces circulating tumor-immune cell clusters and suppresses TNBC metastasis (12). Consistent with their functional role in ICB resistance, tumors expressing high levels of PLXNB1 or PLXNB2 exhibited poorer PFS following anti-PD-1 therapy (Fig. 3E, left). In breast cancer, elevated PLXNB2 predicted worse outcomes in lymph node-positive and TNBC patient subsets (Fig. 3E, right). Mechanistically, ChIP-seq analyses revealed increased co-occupancy of H3K9me3 and H3K36me3, histone marks associated with transcriptional activation at gene bodies (7), across the *PlXNB2* locus in mdig KO cells. These findings suggest that loss of mdig promotes transcriptional activation of inhibitory immune checkpoint genes, potentially contributing to impaired ICB responsiveness.

**Figure 3.**
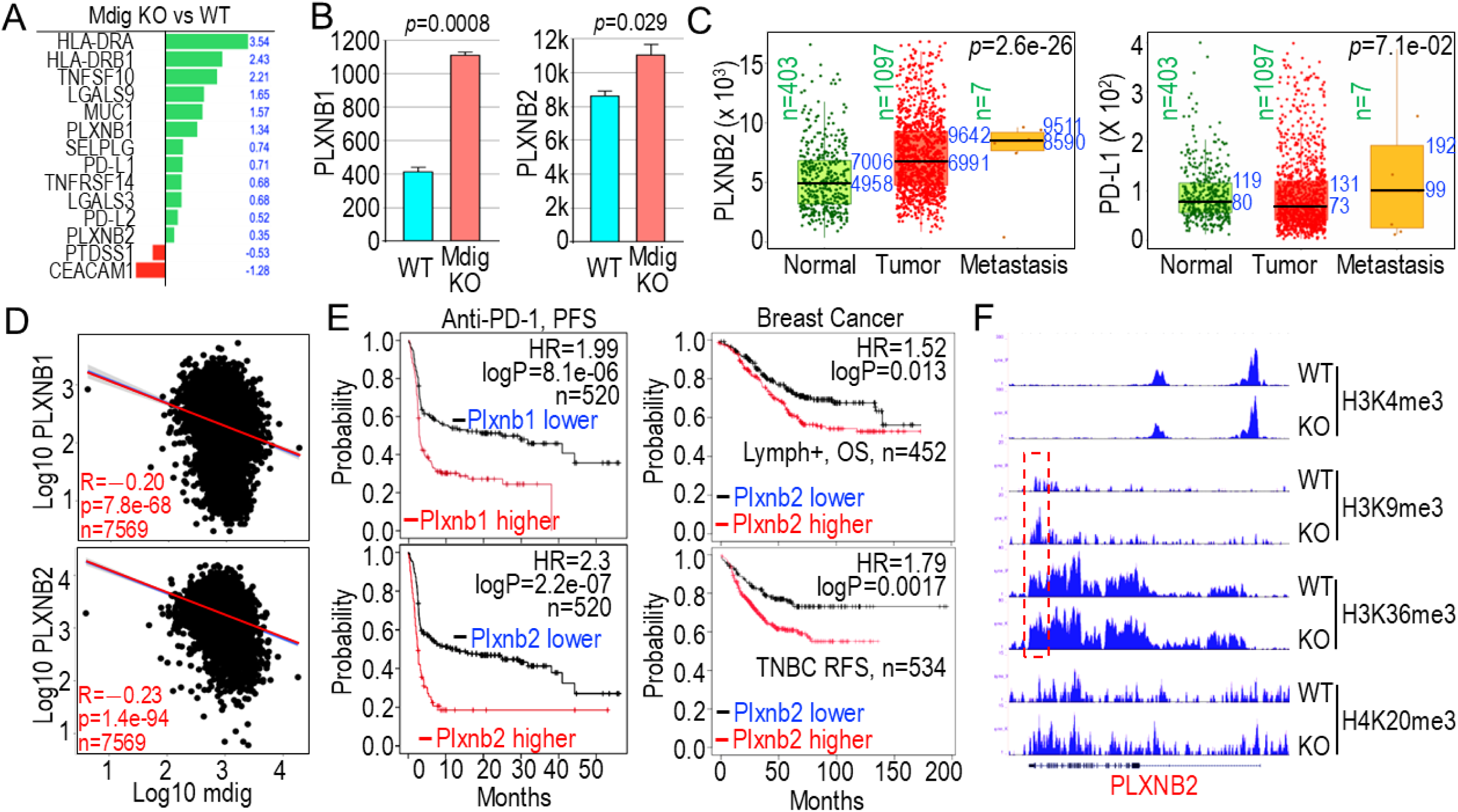
Negative regulation of inhibitory immune checkpoint genes by mdig. **(A, B)** Expression analysis of canonical and non-canonical inhibitory immune checkpoint genes in WT and mdig KO MDA-MB-231 cells. Loss of mdig markedly upregulated multiple checkpoint genes, including *HLA-DRA/DRB1*, *LGALS9*, *PD-L1*, *PD-L2*, *SELPLG*, *PLXNB1*, and *PLXNB2*. **(C)** Elevated PLXNB2 and PD-L1 levels are observed in metastatic lesions compared with normal and primary tumor tissues in breast cancer. **(D)** Clinical correlation analysis showing an inverse relationship between mdig expression and PLXNB1/PLXNB2 levels in patient tumors across multiple cancer types. **(E)** Kaplan-Meier survival analysis showing that tumors with high PLXNB1 or PLXNB2 expression exhibit poorer PFS following anti–PD-1 therapy (left panel). In breast cancer, high PLXNB2 predicts worse outcomes in lymph node-positive and TNBC patient subsets (right panel). **(F)** ChIP-seq analysis of the *PLXNB2* locus in mdig KO cells demonstrating increased co-occupancy of H3K9me3 and H3K36me3, histone marks associated with transcriptional activation.

### Knocking out mdig re-shapes 3D chromatin architecture of immune checkpoint loci

The findings above suggest that mdig may coordinate immune-checkpoint regulation with metastatic behavior in TNBC by restraining the expression of inhibitory immune-checkpoint genes. Mdig has previously been implicated in modulating the deposition of repressive histone methylation marks, including H3K9me3 (13), H3K27me3 (14), and H3K36me3 (5), thereby influencing higher-order chromatin folding and transcriptional output. To delineate how mdig governs 3D genome architecture at immune-checkpoint loci, we performed Hi-C profiling with a focus on *LGALS9, PD-L1, PD-L2*, and *PLXNB2*. TAD and loop calling using Juicebox revealed a pronounced repositioning of both the TAD boundary and major chromatin loops surrounding the *LGALS9* locus in mdig-deficient cells (Fig. 4A), indicating substantial remodeling of local cis-regulatory topology. At the *PD-L1/PD-L2* region, WT cells exhibited a prominent chromatin loop that juxtaposed these genes with an H4K20me3-enriched repressive compartment. This long-range loop was largely abolished upon mdig depletion, potentially relieving structural repression and permitting elevated *PD-L1* and *PD-L2* transcription (Fig. 4B). Remarkably, mdig loss also disrupted the hierarchical TAD and sub-TAD organization across a broader genomic interval containing *HDAC10*, *MAPK12*, *MAPK11*, *PLXNB2*, *PPP6R2*, and *CHKB*, including the collapse of an internal loop (Fig. 4C). This architectural breakdown spatially segregated the H4K20me3-marked repressive domain from *PLXNB2* and neighboring genes, a configuration consistent with the elevated *PLXNB2* expression observed in mdig KO cells. Together, these results demonstrate that mdig maintains repressive chromatin topology at immune-checkpoint and metastasis-associated loci by preserving TAD integrity and long-range loop formation. Loss of mdig dismantles these structures, thereby derepressing immune-evasion gene networks.

**Figure 4.**
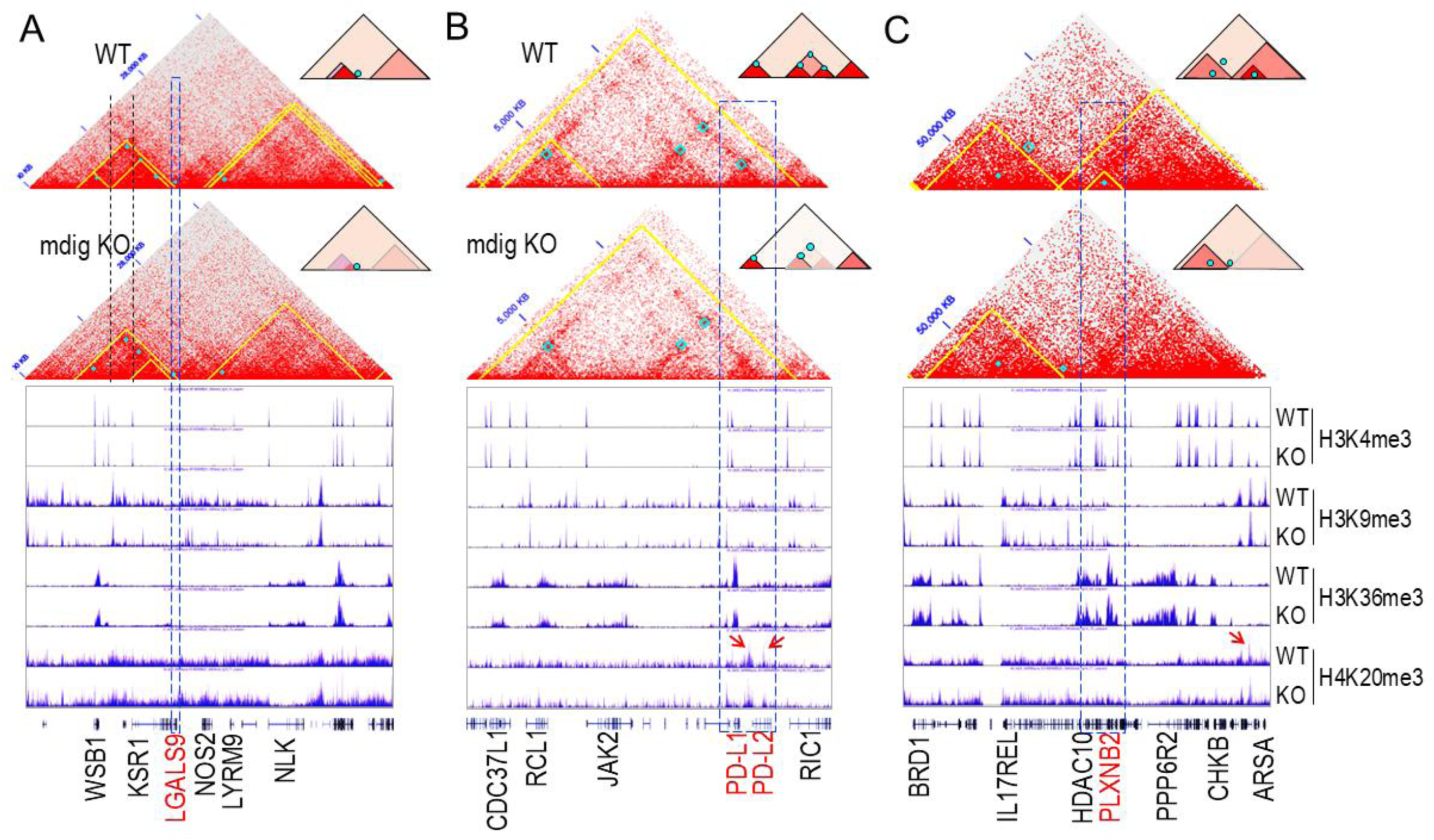
Mdig loss alters 3D chromatin architecture at immune-checkpoint loci. **(A)** Hi-C maps of the *LGALS9* region showing shifts in the TAD boundary and major chromatin loops in mdig-KO cells. **(B)** At the *PD-L1/PD-L2* locus, a long-range repressive loop present in WT cells is abolished upon mdig depletion, consistent with increased gene expression. **(C)** Mdig knockout disrupts TAD/sub-TAD organization across a broader interval containing *HDAC10*, *MAPK12*, *MAPK11*, *PLXNB2*, *PPP6R2*, and *CHKB*, including collapse of an internal loop and separation of an H4K20me3-enriched repressive domain from *PLXNB2*.

### Mdig deficiency causes TAD remodeling through gaining of H3K9me3 and/or H3K36me3

At the whole-chromosome level, several chromosomes, including chromosomes 4, 10, 18, 20, 21 and X, exhibited pronounced alterations in large-scale 3D architecture following mdig knockout. Loss of mdig also increased intra-chromosomal contacts on chromosomes 10, 18 and X (Figs. 5A & 5B, top panel). Our previous work demonstrated that enhanced expression of X-linked genes in mdig-deficient cells was associated with elevated deposition of H3K9me3 and H3K36me3 (7). Consistent with this, chromosome X displayed the most extensive remodeling of TADs and chromatin loops. Integration of Hi-C and ChIP-seq analyses revealed that regions undergoing 3D chromatin reorganization upon mdig depletion were more likely to acquire increased H3K9me3 and/or H3K36me3 (Fig. 5B). Incorporating RNA-seq data further indicated that genes residing within TADs that gained H3K9me3 alone tended to be downregulated in mdig KO cells (Fig. 5B, left). In contrast, genes located in TADs that acquired both H3K9me3 and H3K36me3 were more frequently upregulated, exemplified by a metastatic gene cluster on chromosome X containing *MAGED4*, *SMC1A*, *IQSEC2*, *HUWE1*, and *MAGED2* (Fig. 5B, right). Globally, mdig depletion resulted in a reduced number of chromatin loops (Fig. 5C), a pattern likely reflecting the widespread increase in the repressive histone mark H3K9me3 in mdig KO cells. Together, these findings demonstrate that mdig is a critical guardian of higher-order chromatin organization. Loss of mdig triggers widespread TAD remodeling and loop destabilization driven by aberrant accumulation of H3K9me3 and H3K36me3, leading to mis-regulation of both repressed and activated gene clusters, including immune-checkpoint and metastasis-associated loci. These results position mdig as a key architectural regulator whose deficiency reprograms the 3D genome to favor tumor immune evasion and metastatic progression.

**Figure 5.**
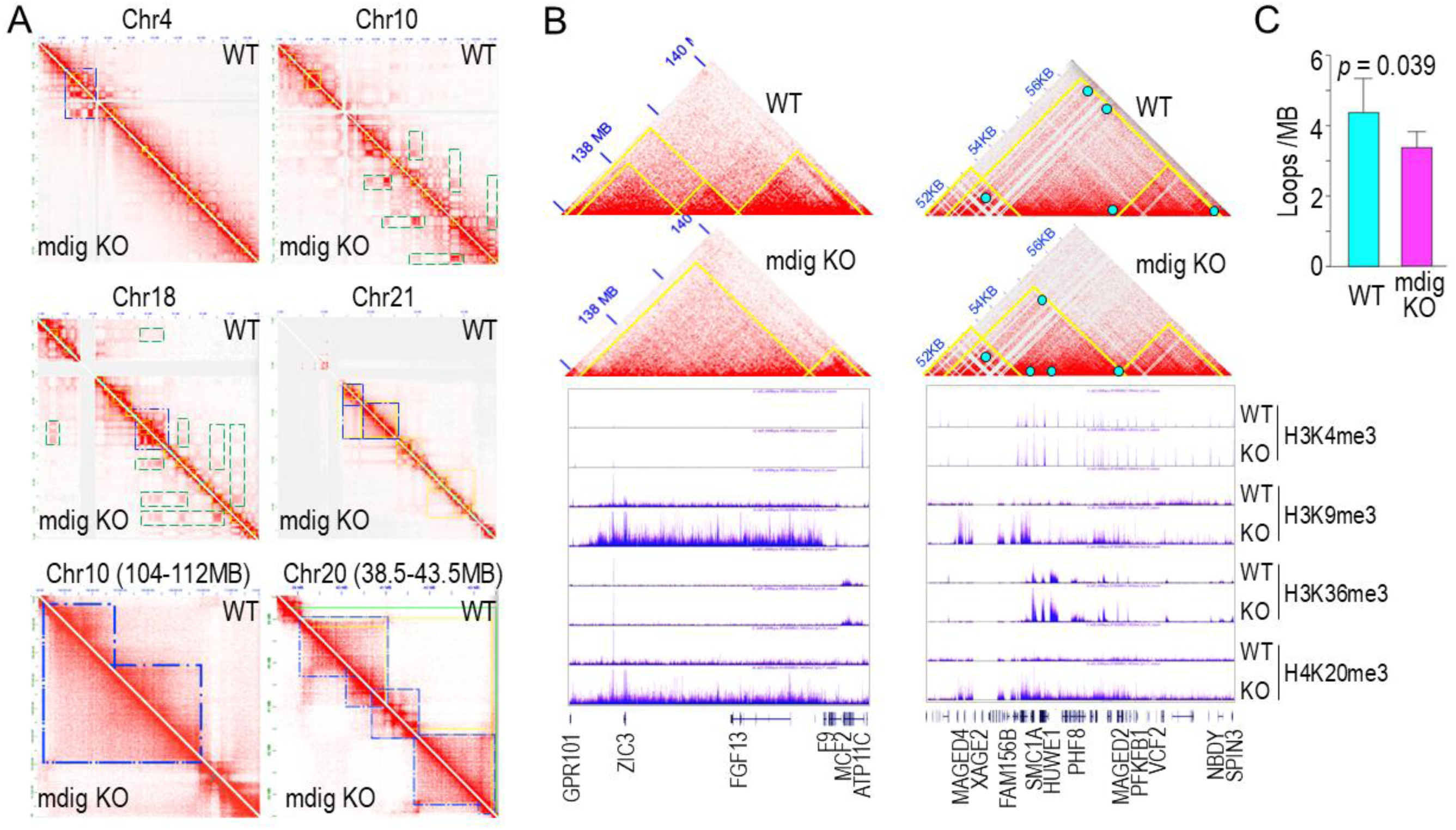
Mdig deficiency induces TAD remodeling and loop destabilization through aberrant gains of H3K9me3 and H3K36me3. **(A)** Genome-wide Hi-C analysis showing large-scale 3D architectural changes across multiple chromosomes (4, 10, 18, 20, and 21) following mdig knockout. Mdig loss increases intra-chromosomal contacts, particularly on chromosomes 10, and 18. **(B)** Hi-C mapping exhibiting the most extensive TAD and loop reorganization on chromosome X following mdig depletion. Integration of Hi-C with ChIP-seq reveals that regions undergoing structural remodeling preferentially gain H3K9me3 and/or H3K36me3. RNA-seq integration shows that genes within TADs gaining H3K9me3 alone are generally downregulated (left), whereas genes in TADs acquiring both H3K9me3 and H3K36me3 tend to be upregulated, exemplified by a metastatic gene cluster on chromosome X (*MAGED4*, *SMC1A*, *IQSEC2*, *HUWE1*, *MAGED2*) (right). **(C)** Mdig knockout reduces the global number of chromatin loops.

## Discussion

Metastasis remains the primary cause of cancer-related mortality and is frequently refractory to conventional and adjuvant therapies [15]. Among breast cancer subtypes, TNBC is associated with the poorest prognosis, driven predominantly by its aggressive metastatic propensity compared with non-TNBC subtypes [16,17]. We previously demonstrated that mdig functions as a suppressor of metastasis, as its genetic ablation promotes EMT and markedly enhances TNBC metastasis *in vivo* [18]. This anti-metastatic role is further substantiated by the current findings: re-expression of mdig in knockout cells significantly attenuates liver metastasis, whereas mdig deficiency upregulates gene networks governing EMT and inhibitory immune checkpoints. Chromatin conformation analysis via Hi-C revealed profound TAD reorganization in mdig KO cells relative to WT counterparts. These architectural alterations strongly correlate with increased deposition of repressive H3K9me3 and active H3K36me3 marks, particularly at loci harboring pro-metastatic and immune-checkpoint genes.

Emerging evidence suggests that the regulatory functions of mdig in Th17 and Treg (8), as well as in IL-4-dependent T-cell programs (15), mechanistically converge with immune checkpoint pathways through the coordinated control of T-cell differentiation, cytokine signaling, and the immunosuppressive tumor microenvironment. In inflammatory and fibrotic settings, mdig deficiency reduces Th17 infiltration and cytokine output while enhancing the recruitment of Foxp3^+^ Tregs, indicating that mdig normally favors a pro-inflammatory Th17 phenotype while suppressing Treg-mediated immune tolerance (16). This balance is mechanistically relevant to checkpoint biology, as Tregs are one of the dominant cellular drivers of immune checkpoint-mediated suppression: they constitutively express high levels of CTLA-4, upregulate PD-1 in suppressive states, and induce PD-L1 expression on antigen-presenting cells through IL-10 and TGF-β that favors exhaustion of the CD8^+^ T cells (9, 17). In cancer patients with relatively high mdig expression, ICBs targeting CTLA-4, PD-1, and PD-L1 exhibited better therapeutic efficacy compared with patients with lower mdig expression, which is uncommon in clinical practice. Consequently, fluctuations in the Th17/Treg balance directly influence checkpoint engagement within the tumor microenvironment, accounting for the worse outcomes and reduced responsiveness to ICB observed in tumors enriched with Tregs (9, 18). However, how the low expression of mdig in tumors contributes to alterations in the tumor microenvironment remains unclear, and the underlying mechanisms require further investigation.

Moreover, by suppressing IL-4 transcription through promoter-level chromatin remodeling, mdig is functionally linked to Th2 polarization (19), a process that indirectly shapes immune checkpoints responses. IL-4 and downstream type-2 cytokines drive alternative macrophage activation (M2 polarization), a state strongly associated with elevated PD-L1, PD-L2, and other immune-inhibitory ligands. IL-4–mediated signaling can also suppress cytotoxic CD8^+^ T-cell activity and promote Treg expansion, further amplifying checkpoint-dependent immunosuppression. Through its effects on chromatin accessibility and transcriptional output, mdig may therefore modulate not only T-cell lineage decisions but also the expression of immune-inhibitory ligands within the tumor microenvironment.

Unlike other breast cancer subtypes, TNBC is enriched for T cell infiltration, including CD8^+^ T cells (20). In TNBC patients who respond to ICB, the increased pool of tumor-infiltrating CD8^+^ cells arises from both clonal reinvigoration of pre-existing CD8^+^ populations and the clonal replacement driven by the influx of newly emerged CD8^+^ clones. Studies in murine TNBC models have revealed two major intratumoral CD8^+^ T cell subsets: tissue-resident memory CD8^+^ cells and terminal exhausted CD8^+^ cells. By repressing immune checkpoint genes and Treg cells, mdig may plausibly shift the balance away from terminal exhaustion and toward functional memory CD8^+^ cells, either through clonal expansion or increased influx of new CD8^+^ clones. Taken together, these mechanisms suggest that mdig governs immune-checkpoint-relevant pathways at multiple levels: (i) by shaping the Th17/Treg axis, which determines the magnitude of CTLA-4- and PD-1–dependent suppression; (ii) by modulating IL-4-driven macrophage and stromal programs that regulate expression of *PD-L1*, *PD-L2*, *LGALS9*, *PLXNB1*, *PLXNB2*, and related genes; and (iii) potentially by altering chromatin architecture at immune-regulatory loci in both tumor and immune cells. Although mdig is not a canonical immune checkpoint molecule, its epigenetic and transcriptional activities position it as an upstream integrator of tumor immune landscape, providing a mechanistic rationale for its strong association with ICB outcomes in TNBC. Clinically, the levels of tumor-derived mdig may serve as a predictive biomarker for identifying the optimal therapeutic window for ICB. By coordinating the Th17/Treg axis, IL-4-dependent programs, and CD8^+^ T cell exhaustion states, mdig captures key immune features that determine responsiveness to immunotherapy.

## Materials and Methods

### Cell culture procedures

MDA-MB-231 triple-negative breast cancer cells (ATCC, cat#CRM-HTB-26) were maintained in DMEM/F-12 medium (GIBCO, cat#10565018) supplemented with 10% fetal bovine serum (GIBCO, cat#A5669701) and 1% penicillin–streptomycin (GIBCO, cat#15140122) at 37 °C in a humidified incubator with 5% CO₂ (Baker ReCO₂ver™).

### Engineering of mdig knockout and overexpression cell lines

To generate mdig-knockout cells, the mdig coding sequence was analyzed using the CRISPR Design Tool (http://crispr.mit.edu/), and an sgRNA targeting exon 3 was selected. The sense and antisense oligonucleotides (5′-CACCGAATGTGTACATAACTCCCGC-3′ and 5′-AAACGCGGGAGTTATGTACACATTC-3′) were annealed at 95 °C for 5 min and cooled to room temperature. The pSpCas9-2A-Blast vector was digested with BbiI (New England Biolabs, cat#R3539S), followed by ligation with the annealed sgRNA duplex using T4 DNA (New England Biolabs, cat#M0202S) ligase at 22 °C for 10 min. The ligation products were transformed into DH5α *E. coli* (Thermo Scientific, cat#EC0112) for plasmid amplification. MDA-MB-231 cells seeded at 2.5 × 10^5^ per well in 6-well plates were transfected with the CRISPR-Cas9 constructs using Lipofectamine 2000 (Invitrogen, cat#11668500) following the manufacturer’s instructions. Forty-eight hours post-transfection, cells were transferred to 10-cm dishes and selected with 2 µg/mL blasticidin (Gibco, cat#A1113903) for two weeks. Individual colonies were screened by Western blotting to identify mdig-knockout (KO) clones, while colonies retaining mdig expression were designated as wild-type (WT).

For mdig overexpression studies, mdig-KO MDA-MB-231 cells were infected with lentiviral particles encoding the MINA53 (NM_153182) Human Tagged ORF (OriGene Technologies, Inc, cat#RC203931L4V and RC203931L3V) along with matched controls. GFP fluorescence was examined 72 h after infection using an EVOS M7000 microscope (Thermo Fisher Scientific). Puromycin-resistant colonies were expanded and validated for Myc-DDK, mdig, and GFP expression by Western blotting.

### Mammary fat pad xenografts of mdig-modified breast cancer cells

All animal procedures were conducted following Stony Brook University DLAR guidelines (IACUC protocol #202100104). Female NSG mice (NOD.Cg-Prkdc^scid^ Il2rg^tm1Wjl^/SzJ; Jackson Laboratories) were housed under controlled conditions with ad libitum access to food and water. MDA-MB-231 WT, mdig KO, vector-control KO, and mdig-reexpressing KO cells were harvested, resuspended in PBS containing phenol-red-free Cultrex BME (10 mg/mL; R&D Systems), and injected (200 μL, 1×10^6^ cells per mouse) into the 5^th^ mammary fat pads of eight-week-old mice. Thirteen mice received WT cells, 14 received mdig KO cells, 6 received vector-control KO cells, 6 received mdig-reexpressing KO cells, and two control mice received PBS. Tumor growth was measured weekly with digital calipers starting 11 days post-transplantation. After nine weeks, mice were euthanized by CO₂ inhalation and perfused with PBS. Primary tumors, lungs, and livers were collected for gross examination and histological evaluation.

### RNA-sequencing and analysis

Total RNA was extracted from 2 × 10^6^ cells using the MagMAX™ mirVana™ Total RNA Isolation Kit (Thermo Fisher, cat#A27828) on a KingFisher Apex system. For each sample, 500 ng of RNA was used to prepare libraries with the Illumina TruSeq Stranded mRNA Library Kit (cat#20020594), followed by paired-end 150-nt sequencing on an Illumina NovaSeq 6000. Reads were aligned to the human GRCh37/hg19 genome, and transcripts were assembled using Cufflinks. Gene expression levels were normalized as geometric Fragments Per Kilobase of transcript per Million mapped reads (FPKMs). Differential expression analysis was performed with DESeq2 (v1.14.1). The RNA-seq dataset has been deposited in NCBI GEO (GSE282125).

### Clinical relevance analysis of mdig in response to ICB

The prognostic impact of mdig (Affymetrix ID: RIOX2) expression in patients receiving immune checkpoint blockade was evaluated using the Kaplan-Meier Plotter platform (http://kmplot.com/analysis/). Survival analyses were performed across immunotherapy cohorts, with stratification by treatment class, including anti-PD-1, anti-PD-L1, and anti-CTLA-4 therapies. The clinical relevance of PLXNB1 and PLXNB2 (probe IDs: 208890_s_at and 211472_at) was assessed in breast cancer datasets independent of ICB datasets. To compare mdig (Affymetrix ID: RIOX2) expression between responders and non-responders to ICB, including pembrolizumab and anti-CTLA-4 therapies, analyses were conducted using the ROC Plotter tool (https://rocplot.com/) across metastatic cancer datasets. Expression patterns of mdig (Affymetrix ID: RIOX2), PLXNB2, and PD-L1 in normal, primary tumor, and metastatic breast tissues were examined using TNMplot (https://tnmplot.com/analysis/). TNMplot correlation analysis was further used to identify genes negatively correlated with mdig in breast cancer.

### Pathway enrichment analysis

Pathway enrichment was conducted using Enrichr (https://maayanlab.cloud/Enrichr/) with KEGG, Reactome, and MSigDB gene set libraries. Differentially expressed genes (DEGs) were defined using fold-change and adjusted p-value thresholds. Genes negatively correlated with mdig were identified through TNMplot correlation analysis (cutoff: Spearman’s Rho ≤ −0.2). Enrichr was subsequently used to calculate enrichment scores and significance values, enabling identification of transcription factors, signaling pathways, and biological processes enriched among DEGs or mdig-negatively correlated gene sets.

### Chromatin immunoprecipitation and sequencing (ChIP-seq)

ChIP-seq analysis was performed following procedures described in our earlier studies (5, 14). All antibodies and reagents were obtained from Active Motif (Carlsbad, CA), and antibody specificity and signal-to-noise ratio were validated prior to use. Genome-wide chromatin states were annotated using histone methylation marks: H3K4me3 (Thermo Fisher, cat#711958), H3K9me3 (CST, cat#4658), H3K36me3 (CST, cat#4909), and H4K20me3 (CST, cat#5737). Data quality was evaluated using mapping efficiency, sequencing depth, normalized strand cross-correlation, background uniformity, and GC bias at peak summits. Enrichment and read distributions were visualized in the UCSC Genome Browser (https://hgw1.soe.ucsc.edu/index.html). The resulting dataset is available at NCBI GEO (GSE207505).

### Hi-C sequencing

Hi-C libraries were generated using the Arima Genomics Hi-C kit following the manufacturer’s protocol. Briefly, cells were crosslinked, nuclei were isolated, and chromatin was digested with the Arima restriction enzyme mix. After end filling with biotin and proximity ligation, crosslinks were reversed and DNA was purified. Biotinylated fragments were enriched, sheared, and converted into Illumina sequencing libraries. Final libraries were quantified, quality-checked, and sequenced on an Illumina platform using paired-end reads. Raw FASTQ files were processed using the Juicer pipeline for read alignment, filtering, and generation of .hic matrices against the hg38 reference genome. Juicebox was used for further visualization. Normalized .hic files generated by the Juicer pipeline were loaded into Juicebox for downstream 3D genome analysis. A/B chromatin compartments were identified based on eigenvector decomposition implemented in Juicebox, using the first principal component. TADs were detected by inspecting insulation patterns and domain boundaries across multiple resolutions.

### Statistical analysis

ChIP-seq and RNA-seq datasets were quality-assessed by Active Motif (Carlsbad, CA), and Hi-C sequencing quality control was performed by Arima Genomics (Carlsbad, CA). All other statistical evaluations were performed using GraphPad Prism 7 (GraphPad Software, San Diego, CA). Experiments were carried out independently at least three times, with a minimum of three biological replicates per condition. Data are expressed as mean ± standard deviation (SD) unless otherwise indicated. Differences between groups were assessed using Student’s t-test or one-way ANOVA, as appropriate. Statistical significance was defined as a two-sided p-value < 0.05.

## Authors’ contributions

**ZW, ZQ**, and **FC** conceived and coordinated the study, analyzed the data and wrote the manuscript. **CT, YQ**, and **MD** performed the cell culture, animal study, Western blotting, and immunohistochemistry; **JW** and **JD** provided technical support.

## Abbreviations

Mdig: mineral dust-induced gene
TNBC: Triple-negative breast cancer
EMT: Epithelial-mesenchymal transition
ICB: Immune checkpoint blockade
TADs: Topologically associating domains
ChIP-seq: Chromatin immunoprecipitation and sequencing
PFS: Progression-free survival
H3K9me3: Histone 3 lysine 9 trimethylation
H3K36me3: Histone H3 lysine 36 trimethylation
H4K20me3: Histone H4 lysine 20 trimethylation
H3K4me3: Histone H3 lysine 4 trimethylation

